# RIPK3 suppresses photoreceptor degeneration in a Stargardt disease mouse model

**DOI:** 10.64898/2026.04.26.720951

**Authors:** Feng Wen, Xiying Zhai, Wanling He, Yaqiong Yang, Liying Tang, Yuehua Zhang, Małgorzata B Różanowska, Xinxuan Cheng, Jiani Li, Jintao Shi, Wei Li, Chunxia Chen, Li Qian, Zhaoqiang Zhang, Yalin Wu, Mouxin Zhang, Jianfeng Wu, Yi Liao

## Abstract

Autosomal recessive Stargardt disease type 1 (STGD1) is the most prevalent inherited juvenile macular degeneration, ultimately leading to irreversible blindness through photoreceptor cell loss, yet the underlying cell death mechanisms remain poorly defined. Receptor-interacting protein kinase 3 (RIPK3) is a well-established mediator of necroptosis and, under certain circumstances, apoptosis downstream of TNF family ligands, and it was found to be progressively upregulated in the retina of a STGD1 *abca4^−/−^rdh8^−/−^* (DKO) mouse model coinciding with the onset and progression of photoreceptor degeneration. Despite elevated RIPK3 expression, necroptosis was not detectable in this model, as evidenced by the absence of phosphorylated MLKL and unaltered photoreceptor degeneration in *abca4^-/-^ rdh8^-/-^mlkl^-/-^* mice. Intriguingly, genetic ablation of *Ripk3* exacerbated photoreceptor loss in DKO mice in both chronic (age-dependent) and acute (light-induced) retinal degeneration paradigms. This detrimental effect was partially ameliorated by pan-caspase inhibition in the acute degeneration model, indicating caspase-dependent apoptosis as the primary executioner. Mechanistically, we demonstrated that RIPK3 suppressed extrinsic apoptosis by attenuating caspase-8 activation downstream of TNF family ligands. Collectively, our findings reveal a non-canonical, protective role of RIPK3 in photoreceptors, as a brake on apoptotic signaling rather than a necroptotic executor, in the context of STGD1. These findings redefine the role of RIPK3 in retinal degeneration and emphasize the contextual plasticity of cell death regulators in neurodegenerative diseases.

## Introduction

Macular degenerative diseases represent a group of currently incurable blinding disorders. Among them, age-related macular degeneration (AMD) is the leading cause of irreversible vision loss in the elderly (1), while autosomal recessive Stargardt disease type 1 (STGD1; OMIM: 248200), caused by mutations in the *ABCA4* gene, is the most common inherited juvenile macular dystrophy (2). Both dry AMD and STGD1 are characterized by progressive, irreversible loss of photoreceptors in the macula (2), which leads to central vision impairment. Despite their clinical significance, the precise molecular mechanisms driving photoreceptor death in these conditions remain incompletely understood, hindering the development of effective neuroprotective therapies.

Dysregulation of visual cycle drives photoreceptor degeneration in STGD1. The ABCA4 protein functions as a flippase that mediates the clearance of all-*trans*-retinal (atRAL), a photoproduct generated upon photoactivation of visual pigments, from the disc membranes of photoreceptor outer segments (2, 3). In the cytoplasm, atRAL is rapidly reduced to all-*trans*-retinol by retinol dehydrogenases, such as RDH8. Notably, pathogenic variants in *RDH8* have also been linked to Stargardt disease (4), underscoring the critical importance of efficient atRAL elimination. Loss-of-function mutations in either *ABCA4* or *RDH8* disrupt this detoxification pathway, leading to pathological accumulation of atRAL. As a reactive aldehyde and potent photosensitizer, excess atRAL directly induces photoreceptor cell death (5, 6). Moreover, it serves as a precursor for cytotoxic bisretinoids, including *N*-retinylidene-*N*-retinylethanolamine (A2E) (7, 8) and retinal dimer (RALdi) conjugates (9). These compounds accumulate as lipofuscin fluorophores in retinal pigment epithelium (RPE), and are widely recognized as key contributors to the pathogenesis of both STGD1 and dry AMD (10, 11). Thus, impaired clearance of atRAL not only causes direct photoreceptor toxicity but also fuels the formation of secondary toxic metabolites, positioning atRAL overaccumulation as a central driver of retinal degeneration in STGD1.

Cytokines of the tumor necrosis factor (TNF) family act as master regulators of both apoptotic and necroptotic cell death, and have been implicated in the pathogenesis of multiple retinal degenerative diseases ^(12)^. Upon ligand binding (e.g., TNFα, FasL, TRAIL), death receptors activate caspase-8–dependent extrinsic apoptosis (13). However, when caspase activity is compromised, signaling can divert toward necroptosis via receptor-interacting protein kinase 3 (RIPK3)–mediated phosphorylation of mixed lineage kinase domain–like pseudokinase (MLKL), culminating in plasma membrane rupture and necrosis (13). In the context of STGD1, TNF signaling appears to be aberrantly activated: *abca4^−/−^rdh8^−/−^* (DKO) mice exhibit marked upregulation of TNFR1 during light-induced photoreceptor degeneration (14, 15), and pharmacological interventions that confer photoreceptor protection reduce TNFR1 expression (15). Nevertheless, whether and how TNF family ligand-driven cell death machinery participates in photoreceptor degeneration in STGD1 remains unclear.

RIPK3 is widely known as a core executor of necroptosis downstream of TNF family ligands, wherein its RIP homotypic interaction motif (RHIM) domain–mediated oligomerization with RIPK1 activates its kinase activity, and subsequently triggers MLKL phosphorylation ^(16-18)^. However, RIPK3 also exhibits context-dependent roles in apoptosis. While early studies described RIPK3 as pro-apoptotic ^(19)^, later work showed that inhibition of the kinase activity of RIPK3 (e.g., with high concentration of GSK’843 or GSK’872) can paradoxically induce apoptosis in vitro ^(20, 21)^. Consistently, mice expressing a kinase-dead RIPK3 (D161N) die embryonically due to excessive RIPK1- and caspase-8-dependent apoptosis (21), revealing an essential anti-apoptotic function of RIPK3 during development. In the retina, RIPK3 has been consistently implicated in photoreceptor degeneration across multiple experimental models. It mediates necroptosis in response to diverse stressors, including retinal detachment (22), dsRNA (23), ocular murine cytomegalovirus infection (24), DNA damage (25), light exposure (26-28), hormonal cues (29), and specifically cone-specific death in *rd10* mice (30). However, a recent study in the STGD1 mouse model *abca4^−/−^rdh8^−/−^*(DKO) mice failed to detect MLKL phosphorylation during light-induced photoreceptor degeneration (31), suggesting RIPK3 may function as a regulator of photoreceptor cell death independent of necroptosis in STGD1.

Here, we demonstrated that the TNF signaling pathway was activated in the retinas of DKO mice following light exposure. However, apoptosis rather than necroptosis was identified as the predominant mode of photoreceptor cell death in both in vivo and in vitro STGD1 models. Unexpectedly, despite its upregulation, RIPK3 does not promote cell death; instead, it functions as a brake on apoptosis. Genetic ablation of *Ripk3* exacerbated retinal degeneration in DKO mice. Phenotypic characterization combined with RNA sequencing revealed that RIPK3 acted as a modulator of TNF signaling to suppress apoptosis in the context of STGD1. Consistently, pan-caspase inhibitor Z-VAD-FMK partially protected photoreceptors in *abca4^−/−^rdh8^−/−^ripk3^−/−^*(TKOR) mice after light challenge. Taken together, our findings uncover an anti-apoptotic role of RIPK3 in photoreceptor cell death. This expands the functional repertoire of RIPK3 in the retinas and provides new insights into the regulatory networks governing photoreceptor survival, with potential implications for therapeutic strategies in STGD1 and potentially dry AMD.

## Methods

### Reagents

The following reagents were obtained commercially: Rabbit anti-RIPK1 antibody (Cell Signaling Technology, Cat.No. 3493, USA, RRID: AB_2305314), Rabbit anti-phospho-RIPK1(S166) antibody (Argio, Cat.No. ARG66476, China, RRID: AB_2861229), Rabbit anti-RIPK3 antibody (Prosci, Cat.No.2283, USA, RRID: AB_203256), Rabbit anti-phospho-RIPK3 (Thr231/Ser232) mAb (Cell Signaling Technology, Cat. No.91702, USA, RRID: AB_2937060), Rabbit anti-MLKL antibody (Abclonal, Cat.No. A19685, China, RRID: AB_2862734), Rabbit anti-phospho-MLKL (Ser345) mAb (Cell Signaling Technology, Cat. No.37333, USA, RRID: AB_2799112), Rabbit anti-TNFα antibody (Abclonal, Cat. No. A20851, China, RRID: AB_3669044), Rabbit anti-cleaved PARP antibody (Cell Signaling Technology, Cat. No. 5625T, USA, RRID: AB_10699459), Mouse anti-PAR antibody (R&D systems, Cat. No.4335-MC-100, USA, RRID: AB_3644339), Rabbit anti-AIF antibody (Abclonal, Cat. No. A0811, China, RRID: AB_2757407), Rabbit anti-GAPDH antibody (Abclonal, Cat.No. A19056, China, RRID: AB_2862549), Rabbit anti-Vinculin antibody (Proteintech, Cat.No. 26520-1-AP, USA, RRID: AB_2868558), all-*trans*-retinal(atRAL) (Sigma, Cat. No. R2500, USA), propidium Iodide (Sigma, Cat. No. P4170, USA).

### Animals and light exposure

*Abca4^-/-^rdh8^-/-^*(DKO) mice were a kind gift from the Krzysztof Palczewski Laboratory from the Department of Pharmacology, Case Western Reserve University, United States. *Ripk3^-/-^*mice and *mlkl^-/-^* mice were kindly provided by Prof. Jiahuai Han’s Laboratory (School of Life Science, Xiamen University) (32). C57BL/6J Wildtype (WT) mice were purchased from the laboratory animal research center, Xiamen University. *abca4^-/-^rdh8^-/-^ripk3^-/-^*(TKOR) and *abca4^-/-^rdh8^-/-^mlkl^-/-^*(TKOM) mice were generated by crossing DKO mice with either *ripk3^-/-^* mice or *mlkl^-/-^*mice, respectively. All the mice used in these studies were backcrossed to have C57BL/6J background without *rd8* mutations (**Figures** S2**A** and **B**) and carried a point mutation to result in Leucine at amino acid residue 450 in RPE65 (**Figure** S2**C**). The PCR DNA fragments around residue 450 were amplified from DNA of mouse tails as described previously (33), and Sanger sequencing was performed using the forward primer subsequently. Mice genotyping primers were provided in **Table** S1. All mice were housed in the animal facility of Xiamen University before and during the experiments and were acclimated to a light schedule with alternating Light Emitting Diode (LED) light (15∼20 lux) and dark for 12 h, and were allowed ad libitum to food and water. For light exposure experiments, 4-week WT, DKO, TKOR or TKOM mice were placed in a dark room strictly protected from light for at least 1 d, and then randomly divided into the following groups: 0 d (control group without mydriasis or light exposure),1 d, 2 d, and 3 d (full pupil dilation with 0.5% tropicamide and exposure to white LED light with an intensity of 10,000 lux (white light lamp, color temperature 6,400K, 4,500 Lumens, model E40-large4U-65W, Yijinrui, Changzhou, China) for 30 min, and then were strictly protected from light until sacrifice (mice number for each group n ≥ 3). Mice were sacrificed at the corresponding time, and the eyeballs or neuroretinas were collected for subsequent experiments. This study was approved by the Animal Ethics Committee of Xiamen University and the Animal Ethics Committee of Xiamen University School of Medicine (XMULAC20200076). Animal experiments were performed following the guidelines of the ARVO Statement for the Use of Animals in Ophthalmic and Vision Research.

### Histological analyses

The histological procedures employed were performed according to previous descriptions with slight modifications (34). The eyeballs of the mice were removed with ophthalmic curved tweezers and placed in FAS eyeball fixative (Servicebio, G1109, China) for 24 h. The eyeballs were taken out, trimmed, and washed with 1× PBS (phosphate buffered saline) solution for 10 min with 3 repeats on a rapid speed orbital shaker. The residual solution on the surface of the eyeball was removed with absorbent paper, put it into a paraffin embedding box, for subsequent dehydration and paraffin embedding. The embedded eyeballs were cut into 6∼8 μm retinal sections. Sections were dewaxed and stained with hematoxylin for 3 min and eosin for 2 min. Samples were observed under an optical microscope (Leica Microsystems) and imaged using a slide scanner.

To quantify the changes in the retina after light exposure, either the number of rows of photoreceptor nuclei or the thickness of the outer nuclear layer (ONL) at 500 μm from the optic nerve head was measured. In addition, the thickness of the ONL in the retina of DKO and TKOR mice was measured in the superior-inferior directions at 250 μm from the optic nerve head.

### TUNEL Assay

TUNEL assay was performed on paraffin sections with a commercially available kit (DeadEnd Fluorometric TUNEL System, Cat. No. G3250; Promega, Madison, WI, United States) following the manufacturer’s instructions. The TUNEL-positive cells in the retinal outer nuclear layer (ONL) of 200-μm length were counted in slides from at least 3 different mice, and the percentage of TUNEL-positive cells was calculated by dividing the number of TUNEL-positive nuclei by the total nuclei in the ONL.

### Intraocular propidium iodide (PI) injection

The experiments were performed following previous reports with slight modifications^(22)^. Briefly, 1 μL of propidium iodide (PI, 2 mg/μL) was injected into the vitreous body of mice on day 3 after light exposure. After 2 h, the eyes were enucleated, cut into 10-μm-thick cryosections, air-dried, and fixed in 100% ethanol for 20 min. Hoechst was used to counterstain the nuclei. For positive control (PC), the slides were incubated in 0.2% Triton X-100 for 15 min, and stained with 50 μg/mL PI and Hoechst for another 15 min. The level of PI-positive immunofluorescence in ONL was analyzed using ImageJ software.

### atRAL treatment

4.55 mg atRAL was added into 400 μl dimethylsulfoxide (DMSO) to prepare a 40 mM stock solution. When different concentrations of atRAL were used to treat 661W cells, the stock solution was diluted accordingly with the culture medium.

### Cell lines and cell viability assay

Cone-like 661W cells were purchased from Shanghai Zishi Biotechnology Company, and they were cloned from retinal tumors of a transgenic mouse line (Shanghai, China)^(35)^. Murine BV2 microglia cells were a kind gift from Prof. Xiaofen Chen, Xiamen University. Both cell lines were maintained in Dulbecco’s modified Eagle’s medium (DMEM) media (Gibco, Cat. No. C11995500BT, China) supplemented with 10% fetal bovine serum (Biological Industries, Cat. No. 04-001-1A, Israel) and 1% penicillin/streptomycin (Thermo Fisher Scientific, Cat. No. 15140122, Grand Island, NY, United States) in a humidified incubator with 5% CO_2_ at 37 °C. *Ripk3^-/-^* 661W cells were generated following the protocols published previously (36). The sgRNAs used to generate *ripk3^-/-^* 661W cells were as follows: forward: 5’-CACCGTGGGACTTCGTGTCCGGGC-3’; reverse: 5’-AAACGCCCGGACACGAAGTCCCAC-3’.

To measure cell viability, cells were seeded as 8000 cells/100 μL media/well in a 96-well plate. Cells were treated with different concentrations of atRAL. At 6 h after atRAL treatment, cell counting kit-8 (CCK8, RM02823, Abclonal, China) solution was 1:9 diluted with media, and the mixture was used to incubate cells for another 2 to 3 h in the dark. Cell viabilities were determined by measuring the optical absorbance at 450 nm with a microplate reader (Bio-Tek Instruments, United States).

### RNA extraction and qRT-PCR

The experiments were performed essentially as previously described (37). Two retinas from each mouse without treatment or at 1 d, 2 d, or 3 d after light exposure or cells in 6-well plates were collected into 500 μL Total RNA Extraction Reagent (Trizol, Abclonal, Cat. No. RK30129, Wuhan, China). RNA was extracted according to the manufacturer’s instructions, and diluted in 30∼40 μL DEPC-treated water (Sangon, Cat. No. B501005, Shanghai, China). cDNA was generated using Hifair®Ⅲ 1st Strand cDNA Synthesis SuperMix for qPCR (gDNA digester plus) (Yeasen, Cat. No. 11141ES60, Shanghai, China). Then, for retinal tissues, the mRNA expression levels of *Rik1*, *Casp8*, and *Casp3* were determined by quantitative real-time PCR (qRT-PCR), and *Atp5b* was used as a control. For cell experiments, *Actb* was used as a control. The primers used for qRT-PCR analysis are listed in Table S2.

### Western blotting

Western blot analysis procedures were carried out as formerly descried with slight modifications (37). Mouse neuroretina was dissected and collected in cold RIPA lysis buffer (Pierce, Cat. No. 89900, United States) with 1 × proteinase/phophatase inhibitor cocktail (Apexbio, Cat.No. K1008/K1015, United States). Five to six 1 mm steel beads were added into each tube, and then the neuroretina lysates were ground at 70 Hz for 10 min (45 s on, 15 s off). The lysates were centrifuged at 13,000 rpm for 10 min at 4°C, and supernatants were collected. Equal amounts of proteins (about 30 μg) from each sample were used for Western blotting analysis. Briefly, protein was subjected to electrophoresis in 10% SDS-polyacrylamide gels and transferred to polyvinylidene difluoride membranes (Roche Applied Science, Mannheim, Germany). After blocking with 5% skim milk for 1 h at room temperature, the membranes were incubated with primary antibodies (1:1000 dilution) at 4 °C overnight, followed by incubation with the corresponding secondary antibodies (1:5000 dilution) for 2 h at room temperature. Membranes were developed using ECL Western blotting detection reagents (New Cell & Molecular Biotech Co.Ltd, Cat. No: P10300, Suzhou, Jiangsu, China), and blots were analyzed using a ChemiDoc XRS Imaging system (Bio-Rad, California, USA) and quantified using Quantity One software (Bio-Rad, California, USA).

### Immunoprecipitation

Immunoprecipitation was carried out according to a previously published protocol, with slight adjustments (22). Each retinal lysate sample was prepared in NP-40 lysis buffer (50 mM Tris-HCl pH 7.50, 150 mM NaCl, 1% NP-40, 5% glycerol, 1 mM EDTA, 1 mM EGTA, 10 mM β-glycerophosphate) from 4 mouse retinas. Equal amounts of retinal lysates (500 μg) were incubated with 4 μg anti-RIP1 (BD Biosciences, Cat. No. 610458, USA, RRID: AB_397832) and 20 μL of protein A/G agarose beads (Yeasen, Cat. No. 36403ES03, Shanghai, China) at 4°C overnight. After centrifugation, the protein-antibody-beads complex was washed with lysis buffer 5 times, and elutes were prepared by boiling the complex in 50 μL 1 × SDS loading buffer at 95 °C for 5 min.

### Transmission Electron Microscopy

Mice were sacrificed, and their eyeballs were removed immediately into prechilled 2.5% glutaraldehyde solution for 1 h with a puncture in the cornea. Retina-choroid-sclera complexes were then separated from the anterior segment and vitreous. The central retina-choroid-sclera complexes of the inferior retina about 1 mm from the optic disc were cut into 1 mm^3^ blocks, which were perfused with 2% formaldehyde and 2.5% glutaraldehyde in 0.15 mol/L phosphate buffer, pH 7.4, followed by 1% OsO_4_ and 1.5% potassium ferrocyanide, and stained with 1% uranyl acetate, as previously described^(37)^. The photographs of photoreceptors were obtained with an HT-7800 electron microscope (Hitachi, Tokyo, Japan). Photoreceptors showing nuclear condensation were classified as apoptotic cells, whereas photoreceptors with cellular and organelle swelling were considered to be undergoing necrosis.

### Intraocular administration of pan-caspase inhibitor Z-VAD-FMK or Caspase 8 inhibitor Z-IETD-FMK injection

After full pupil dilation with 0.5% tropicamide, mice were exposed to 10,000 lux of strong white light for 30 min. Afterwards, mice were euthanized. Z-VAD-FMK or Z-IETD-FMK was prepared as 40 μg/μL stock solutions in DMSO, and diluted with 1 × PBS to a final concentration of 4 μg/μL right before injection. 1 μL of Z-VAD-FMK/Z-IETD-FMK was injected into the vitreous body of the treated eyes, whereas 1 μL of DMSO was injected into the control eyes.

### Bulk RNA sequencing, differential expression, and functional enrichment

Retina samples were collected from 3∼4 different mice for each genotype 2 d after light exposure. RNA purification, library preparation, and sequencing were conducted by Majorbio Biotechnology Company, Shanghai, China. RNA-seq data have been deposited in the NCBI Sequence Read Archive (SRA) database (SRP634776). To identify DEGs (differential expression genes) between WT and DKO or DKO and TKOR mouse retinas, the expression level of each transcript was calculated according to the transcripts per million reads (TPM) method. DEGs with |log2FC| > 2 and *p-*value < 0.01 were considered to be significantly different expressed genes for the comparison between DKO and WT mouse retinas, whereas |log2FC| > 1.2 and *p-*value < 0.05 were considered to be significantly different expressed genes for the comparison between TKOR and DKO mouse retinas. In addition, functional enrichment analysis, including GO and KEGG, was performed based on DEGs, which were performed on the online platform of Majorbio Cloud Platform (https://cloud.majorbio.com/).

### Statistical analyses

Statistical analysis was performed using GraphPad Prism software, version 5.0 (http://www.graphpad.com/, RRID: SCR_002798). All data were presented as the mean ± standard deviation (S.D.) with at least three independent biological repeats. The statistical significance of the difference between the experimental and control groups was evaluated by Mann-Whiteney *U* test, one-way, or two-way analysis of variance (ANOVA) followed by *post*-hoc analysis.

## Results

### Activation of TNF Signaling in the Neuroretinas of DKO Mice After Light Exposure

Due to the loss of two genes responsible for the clearance of atRAL, *abca4^-/-^ rdh8^-/-^*(DKO) mice display rapid retinal degeneration following light exposure (34). Accordingly, a light-induced retinal degeneration model was established by exposing DKO mice to 10,000 lux white LED light for 30 min, with wildtype (C57BL/6J, WT) mice serving as experimental controls (**Figure** 1**A**). At 4 weeks of age, without light exposure, the thickness of the outer nuclear layer (ONL) was comparable in WT and DKO mice (**Figures** 1**A** and **B**, *p* = 0.4955, n = 9 for WT Ctrl, n = 9 for DKO Ctrl). After exposure to light, the integrity of WT mouse retina remained morphologically intact (**Figure** 1**A**). In contrast, the average number of nuclei per column in the ONL of the central retina decreased from 12 to approximately 10 at day 1, and further decreased to 7 nuclei at day 3 in DKO mice (*p* < 0.0001, n = 9 for DKO Ctrl, n = 9 for DKO at day 3) (**Figures** 1**A** and **B**).

**Figure 1.**
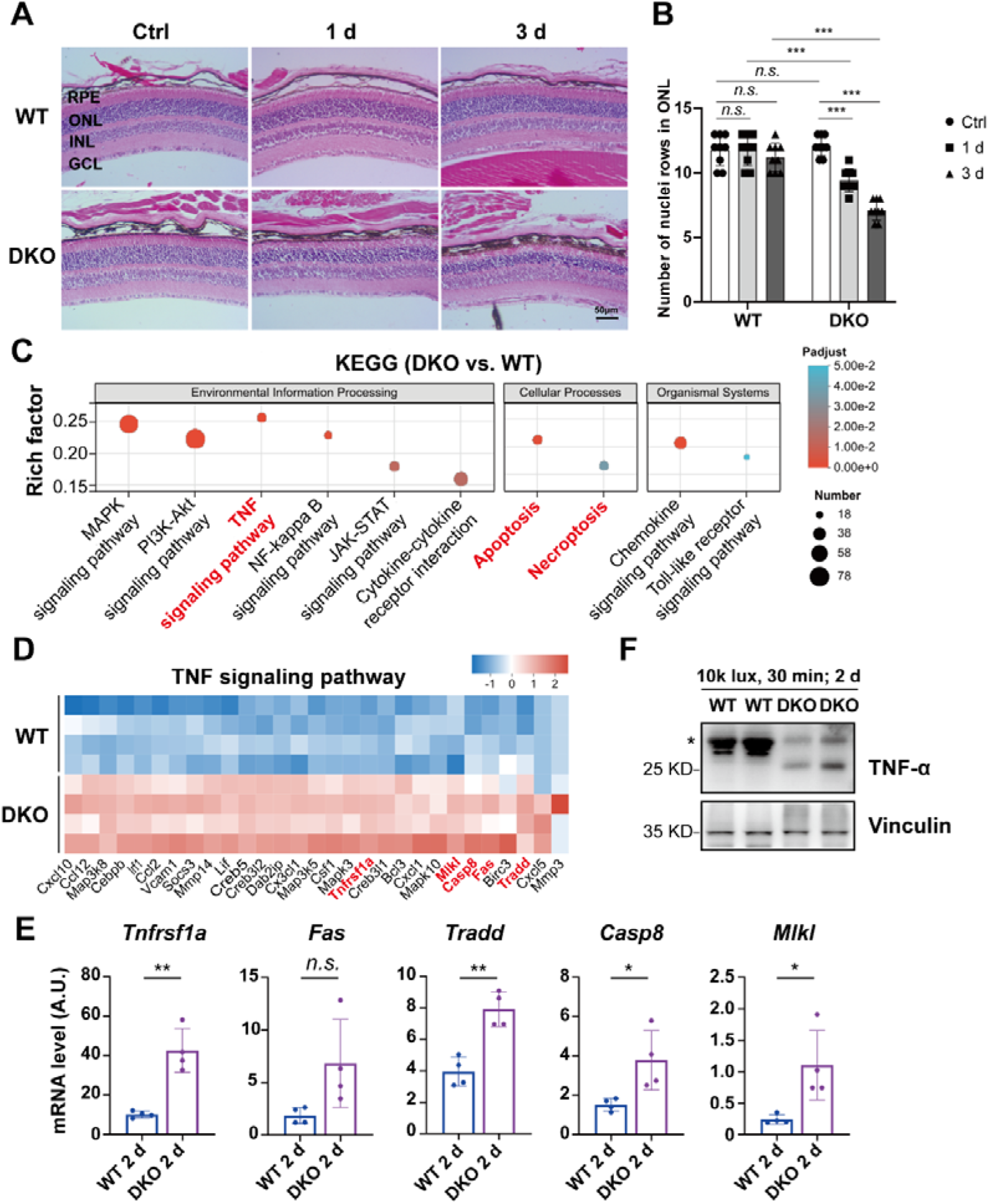
TNF signaling pathway is activated in the retinas of *abca4^-/-^rdh8^-/-^*(DKO) mice after light exposure. (**A**) Representative hematoxylin and eosin (H&E) staining images of mouse retinas from wildtype (WT) and *abca4^-/-^ rdh8^-/-^* (DKO) mice before and after a 30-min exposure to 10,000 lux white LED light (Scale bar = 50 μm). RPE: retinal pigment epithelium; ONL: outer nuclear layer; INL: inner nuclear layer; GCL: ganglion cell layer. (**B**) Quantification of ONL thickness in the central retina of WT and DKO mice. Data were obtained from 9 mice per group. (**C**) KEGG pathway enrichment analysis of differentially expressed genes (DEGs) in the neuroretinas of DKO and WT mice at 2 d after 30-min exposure to white LED light. (**D**) Heatmap of DEGs in TNF signaling pathway in the neuroretinas of WT and DKO mice at 2 d after light exposure. (**E**) Bar graphs of mRNA levels of DEGs in TNF signaling pathway related to apoptosis and necroptosis in the neuroretinas of WT and DKO mice. A.U.: arbitrary units. (**F**) Representative Western blot images of TNFα expression in the neuroretinas of WT and DKO mice at 2 d after light exposure. Vinculin was used as a loading control. For all the experiments, statistical significance was determined by non-parametric Mann-Whitney *U* test or two-way ANOVA with Tukey’s post-hoc test: \**p* < 0.05, \*\**p* < 0.01, \*\*\**p* < 0.001, *n.s.*: not significant. All data are shown as the mean ± standard deviation (S.D.).

To explore the underlying mechanisms regulating photoreceptor degeneration in stress-sensitive DKO mice, neuroretina tissues were collected from WT and DKO mice at 2 d after light exposure for bulk RNA sequencing (RNA-seq) analysis. Using WT retinas as controls, genes with a fold change (FC) ≥ 2 and *p*-value < 0.01 were defined as significantly differentially expressed genes (DEGs). In total, 751 genes were upregulated, whereas 1017 genes were downregulated in DKO retinas compared to WT retinas after light exposure. Kyoto Encyclopedia of Genes and Genomes (KEGG) pathway enrichment results showed that the TNF signaling pathway (mmu04668) was enriched based on upregulated DEGs (**Figure** 1**C**). More interestingly, both the Apoptosis pathway (mmu04210) and Necroptosis pathway (mmu04217)–two downstream cascades that can be triggered by TNF signaling, were also significantly enriched (**Figure** 1**C**). 29 upregulated genes in TNF signaling pathway were visualized in a heatmap (**Figure** 1**D**). Among them, regulators of the canonical extrinsic apoptotic pathway, including *Tnfrsf1a*, *Tradd,* and *Casp8,* were markedly upregulated in light-exposed DKO retinas relative to WT retinas (**Figure** 1**E**, *Tnfrsf1a*: *p* = 0.0012; *Tradd*: *p* = 0.0015; *Casp8*: *p* = 0.0260). Though not statistically significant, the mRNA expression level of *Fas*, another death receptor gene upstream of the extrinsic apoptosis pathway, was also increased in DKO retinas (**Figure** 1**E**, *p* = 0.0584). In addition, we observed a statistically significant upregulation of the key necroptosis regulator gene *Mlkl* in the retinas of light-exposed DKO mice (**Figure** 1**E**, *p* = 0.0215). Moreover, Western blot analysis revealed that mature TNFα proteins were detected in DKO retinas at 2 d after light exposure but not in WT retinas (**Figure** 1**F**). Collectively, these data indicate that TNF signaling can be activated in the retinas of DKO mice following light exposure, which might subsequently trigger apoptosis and necroptosis in photoreceptors.

### Activation of Extrinsic Apoptosis in DKO Mouse Retinas and atRAL-treated 661W Cells

Subsequently, genes that were upregulated in light-exposed DKO mouse retinas and enriched in the Apoptosis pathway were subjected to protein-protein interaction (PPI) network analysis. Within the PPI network, the function importance of a given protein could be directly visualized through the size of its representative node. *Casp8*, which is the major initiator of canonical extrinsic apoptosis pathway, stood as a dominant node in this network (**Figure** 2**A**). Given that multiple death receptor genes, including *Tnfrsf1a*, *Fas*, and *Tnfrsf10b*, were upregulated in DKO retinas, the extrinsic apoptosis pathway might be activated in the retinas of DKO mice after light exposure (**Figure** 2**A**).

**Figure 2.**
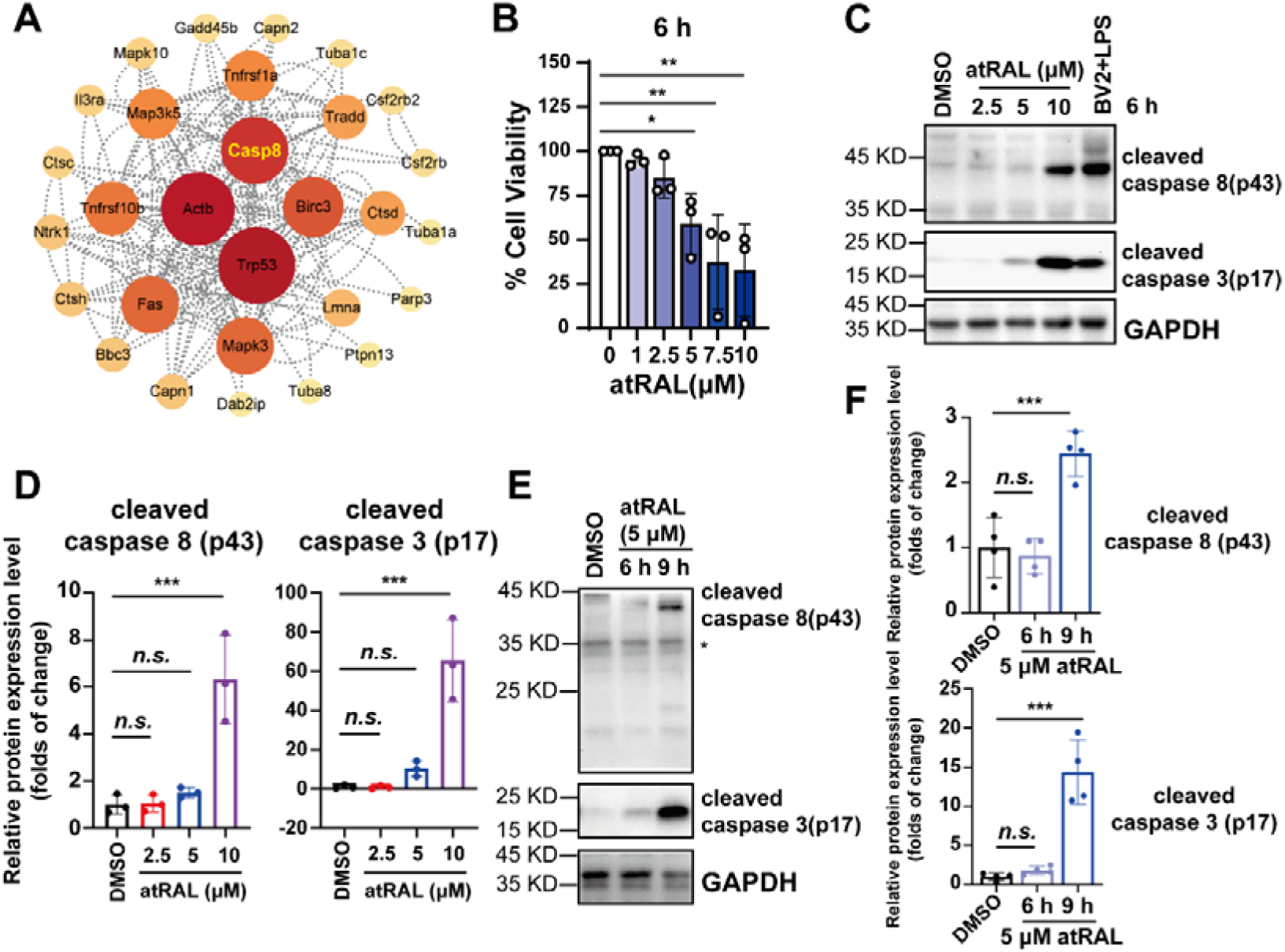
Activation of the extrinsic apoptosis in the photoreceptors after exposure to over-accumulated atRAL. (**A**) Protein-protein interaction network displaying core targets in the enriched apoptosis pathway in the neuroretina of DKO mice after light exposure. (**B**) 661W cells were incubated with increasing concentrations of atRAL for 6 h, and cell viability was evaluated by CCK8 assay. (**C**) Representative Western blot images showing cleaved caspase 8(p43) and cleaved caspase 3(p17) in 661W cell lysates following treatment with increasing concentrations of atRAL for 6 h. GAPDH was used as a loading control. BV2 cells treated with 10 ng/mL LPS were used as a positive control. (**D**) Quantification results of the protein levels of cleaved caspase 8(p43) and cleaved caspase 3(p17) in (**C**). (**E**) Representative Western blot images showing cleaved caspase 8 (p43) and cleaved caspase 3(p17) in 661W cell lysates following treatment with 5 μM atRAL for 6 h and 9 h. GAPDH was used as a loading control. (**F**) Quantification results of the protein levels of cleaved caspase 8 (p43) and cleaved caspase 3 (p17) in (**E**). Data are obtained from 3 or 4 independent experiments. \**p* < 0.05, \*\**p* < 0.01, \*\*\**p* < 0.001, one-way ANOVA with Turkey’s post-hoc analysis. All data are shown as mean ± standard deviation (S.D.).

Rapid accumulation of atRAL after light exposure causes photoreceptor degeneration in DKO mice ^(34)^. Accordingly, an in vitro model was established by using atRAL to treat cone-like 661W cells. Treatment with atRAL reduced the viability of 661W cells in a concentration-dependent manner, with an IC_50_ (half-maximal inhibitory concentration) of 6.11 μM after 6 hours of exposure (**Figure** 2**B**). Western blot analysis revealed the presence of cleavage products of caspase 8(p43) and caspase 3(p17) in atRAL-treated cells when the concentration of atRAL was escalated (**Figures** 2**C** and **D**, caspase 8(p43): DMSO vs. 2.5 μM, *p* > 0.9999; DMSO vs. 5 μM, *p* = 0.8667; DMSO vs. 10 μM, *p* = 0.0004; caspase 3(p17): DMSO vs. 2.5 μM, *p* > 0.9999; DMSO vs. 5 μM, *p* = 0.5962; DMSO vs. 10 μM, *p* = 0.0002).Similarly, when cells were incubated with 5 μM atRAL for 9 h, the cleavage products of caspase 8(p43) and caspase 3(p17) were generated (**Figure** 2**E** and **F**, caspase 8(p43): DMSO vs. 5 μM atRAL 6 h, *p* = 0.8372; DMSO vs. 5 μM atRAL 9 h, *p* = 0.0007; caspase 3(p17): DMSO vs. 5 μM atRAL 6 h, *p* = 0.8528; DMSO vs. 5 μM atRAL 9 h, *p* < 0.0001).Thus, we conclude that extrinsic apoptosis pathway is activated in photoreceptor cells in STGD1 models.

### Necroptosis is not Activated in DKO Mouse Retinas and atRAL-treated 661W Cells

Downstream of TNF signaling, necroptosis is a regulated form of lytic cell death mediated by the sequential phosphorylation of RIPK3 and MLKL ^(38, 39)^. Next, we tried to elucidate whether necroptosis was involved in photoreceptor degeneration in DKO mice. Neuroretina tissues were harvested from DKO mice either from the untreated control group or at 1 d, 2 d, and 3 d after 30-min exposure to 10,000 lux white LED light. While RIPK3, and MLKL were expressed in the retinas, the phosphorylation of these proteins, a post-translational modification specifically required for the induction of necroptosis, was undetectable (**Figure** 3**A**). Notably, however, a statistically significant elevation in the expression of RIPK3 was observed in the retinas of DKO mice after light exposure (**Figure** 3**B**, RIPK3: 0d vs. 1d, *p* = 0.3450; 0d vs. 2d, *p* = 0.0005; 0d vs. 3d, *p* < 0.0001; MLKL: 0d vs. 1d, *p* = 0.2378; 0d vs. 2d, *p* = 0.8670; 0d vs. 3d, *p* = 0.3274). Furthermore, the phosphorylation status of RIPK3 and MLKL was also examined in atRAL-treated 661W cells. Consistent with the results obtained from the neuroretina tissues of DKO mice, the phosphorylated forms of RIPK3 and MLKL remained undetectable in the atRAL-treated cells (**Figures** 3**C** and **D**).

**Figure 3.**
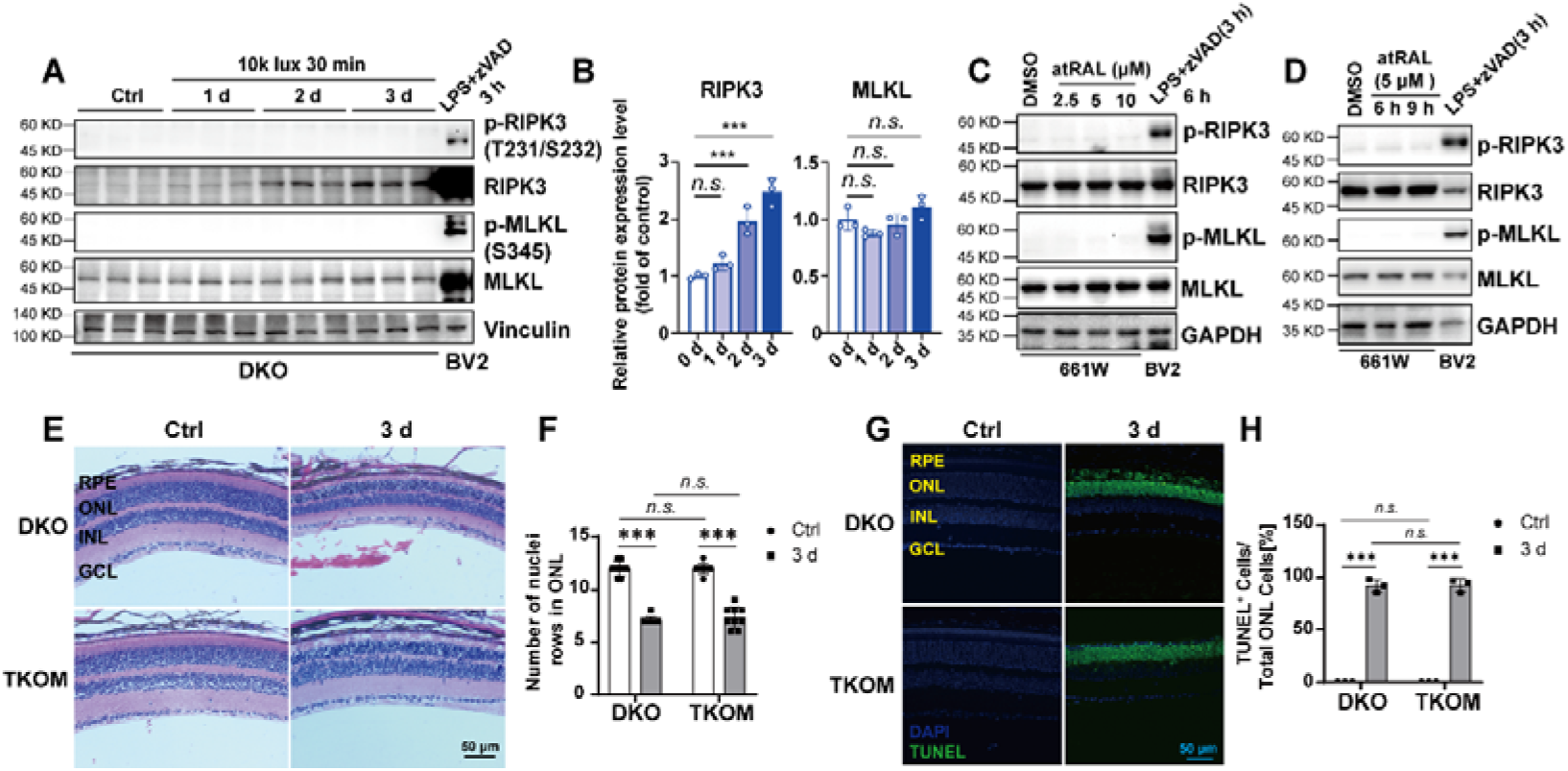
Necroptosis is not involved in photoreceptor degeneration in vivo and in vitro. (**A**) Western blot images of phosphorylation modifications and total proteins of RIPK3 and MLKL in the neuroretinas of DKO mice before and after 30-min exposure to 10,000 lux white LED light. BV2 cells treated with 10 ng/mL Lipopolysaccharide (LPS) plus 50 μM Z-VAD-FMK (zVAD) for 3 h were served as necroptosis positive control. Vinculin was used as a loading control. **(B)** Quantification of the protein levels of RIPK3 and MLKL in (**A**). Data are obtained from 3 mice for each time points. (**C**) Representative Western blot images of phosphorylation and total protein levels of RIPK3 and MLKL in 661W cell lysates after treatment with increasing concentrations of atRAL for 6 h. (**D**) Representative Western blot images of phosphorylation and total protein levels of RIPK3 and MLKL in 661W cell lysates following treatment with 5 μM atRAL for 6 h and 9 h. For (**C**) and (**D**), GAPDH was used as a loading control. BV2 cells treated with 10 ng/mL LPS plus 50 μM zVAD were used as necroptosis positive control. (**E**) Representative hematoxylin and eosin (H&E) staining images of retinal cross-sections from DKO and *abca4^-/-^ rdh8^-/-^ mlkl^-/-^*(TKOM) mice before or at 3 d after 30-min exposure to 10,000 lux white LED light (Scale bar = 50 μm). (**F**) Quantification of ONL thickness in the central retina of DKO and TKOM mice. Data were collected from 9 mice per group. (**G**) Representative terminal deoxynucleotidyl transferase dUTP nick end labeling (TUNEL) staining images of retinal cross-sections from DKO and TKOM mice before or at 3 d after 30-min exposure to 10,000 lux light (Scale bar = 50 μm). (**H**) Quantification of the percentage of TUNEL-positive cells in the ONL of DKO and TKOM mice. Data are obtained from 3 mice per group. RPE: retinal pigment epithelium; ONL: outer nuclear layer; INL: inner nuclear layer; GCL: ganglion cell layer. For all the experiments, statistical significance was determined by one-way or two-way ANOVA with Tukey’s post-hoc test: \**p* < 0.05, \*\**p* < 0.01, \*\*\**p* < 0.001, *n.s.*: not significant. All data are shown as the mean ± standard deviation (S.D.).

As the terminal executor of the necroptosis pathway, MLKL oligomerizes upon activation to permeabilize the membrane lipid bilayer and induce cell death (32, 40, 41). To further verify whether necroptosis is involved in light-induced retinal degeneration in DKO mice, *abca4^-/-^ rdh8^-/-^ mlkl^-/-^* (TKOM) mice were generated. At 3 d after light exposure, the number of nuclei per column in the ONL of central retina was reduced to ∼7 in both DKO and TKOM mice (**Figure** 3**E**, DKO Ctrl vs. DKO 3d, *p* < 0.0001; TKOM Ctrl vs. TKOM 3d, *p* < 0.0001; DKO Ctrl vs. TKOM Ctrl, *p*>0.9999; DKO 3d vs. TKOM 3d, *p* = 0.8997). In addition, TUNEL staining of retinal cross-sections revealed massive cell death in the ONL, and no difference in the percentages of TUNEL-positive cells between DKO and TKOM mice was detected (**Figure** 3**F**, DKO Ctrl vs. DKO 3d, *p* < 0.0001; TKOM Ctrl vs. TKOM 3d, *p* < 0.0001; DKO Ctrl vs. TKOM Ctrl, *p* > 0.9999; DKO 3d vs. TKOM 3d, *p* = 0.9996). Taken together, these results suggest that necroptosis is not involved in photoreceptor degeneration in STGD1 models.

### Knockout of *Ripk3* Exacerbated Photoreceptor Degeneration in DKO Mice

Although our data indicated that necroptosis was not involved in photoreceptor degeneration in DKO mice, it is intriguing to observe that the protein levels of RIPK3 were elevated during photoreceptor degeneration (**Figure** 3**A**). To investigate whether RIPK3 exerted a regulatory role in photoreceptor degeneration in DKO mice, *abca4^-/-^rdh8^-/-^ ripk3^-/-^* (TKOR) mice were generated (**Figure** 4**A**). Because atRAL clearance is impaired ^(42)^, DKO mice are not only susceptible to light-induced retinal injury, but also develop age-dependent photoreceptor dystrophy ^(34, 43)^. Surprisingly, TKOR mice started to show a reduction in ONL thickness at 6 months of age, with further progression of this thinning by 8 months (**Figure** 4**B**). Consistent with the observations in age-dependent models, while loss of *Ripk3* alone did not induce retinal degeneration (**Figure** S1), genetic ablation of *Ripk3* in DKO mice exacerbated photoreceptor degeneration after light exposure (**Figures** 4**C** and **D**). Starting from day 2 after light exposure, TKOR mice exhibited significantly more severe photoreceptor degeneration in the inferior retina compared with DKO mice (**Figure** 4**D**, *p* < 0.0001; 500 μm, *p* = 0.0007; 750 μm, *p* = 0.0007). TUNEL staining and quantification results revealed a significantly greater number of TUNEL-positive cells in the ONL of TKOR mice relative to DKO mice at 2 d after light exposure (**Figures** 4**E** and **F**, DKO Ctrl vs. DKO 2d, *p* = 0.7201; TKOR Ctrl vs. TKOR 2d, *p* = 0.0001; DKO Ctrl vs. TKOR Ctrl, *p* > 0.9999; DKO 2d vs. TKOR 2d, *p* = 0.0001). Moreover, immunostaining results revealed that the expression levels of rod-specific rhodopsin and cone-specific arrestin were markedly lower in TKOR mice compared to DKO mice at 3 d post-light exposure (**Figure** 4**G**), indicating that *Ripk3* deficiency exacerbated the degeneration of both rod and cone photoreceptors. Genotyping results confirmed that the exacerbation of retinal degeneration in TKOR mice was not caused by the Rd8 mutations or mutation in *Rpe65* gene (**Figure** S2). All these results suggest that deletion of *Ripk3* exacerbates photoreceptor degeneration in mice lacking both *Abca4* and *Rdh8*.

**Figure 4.**
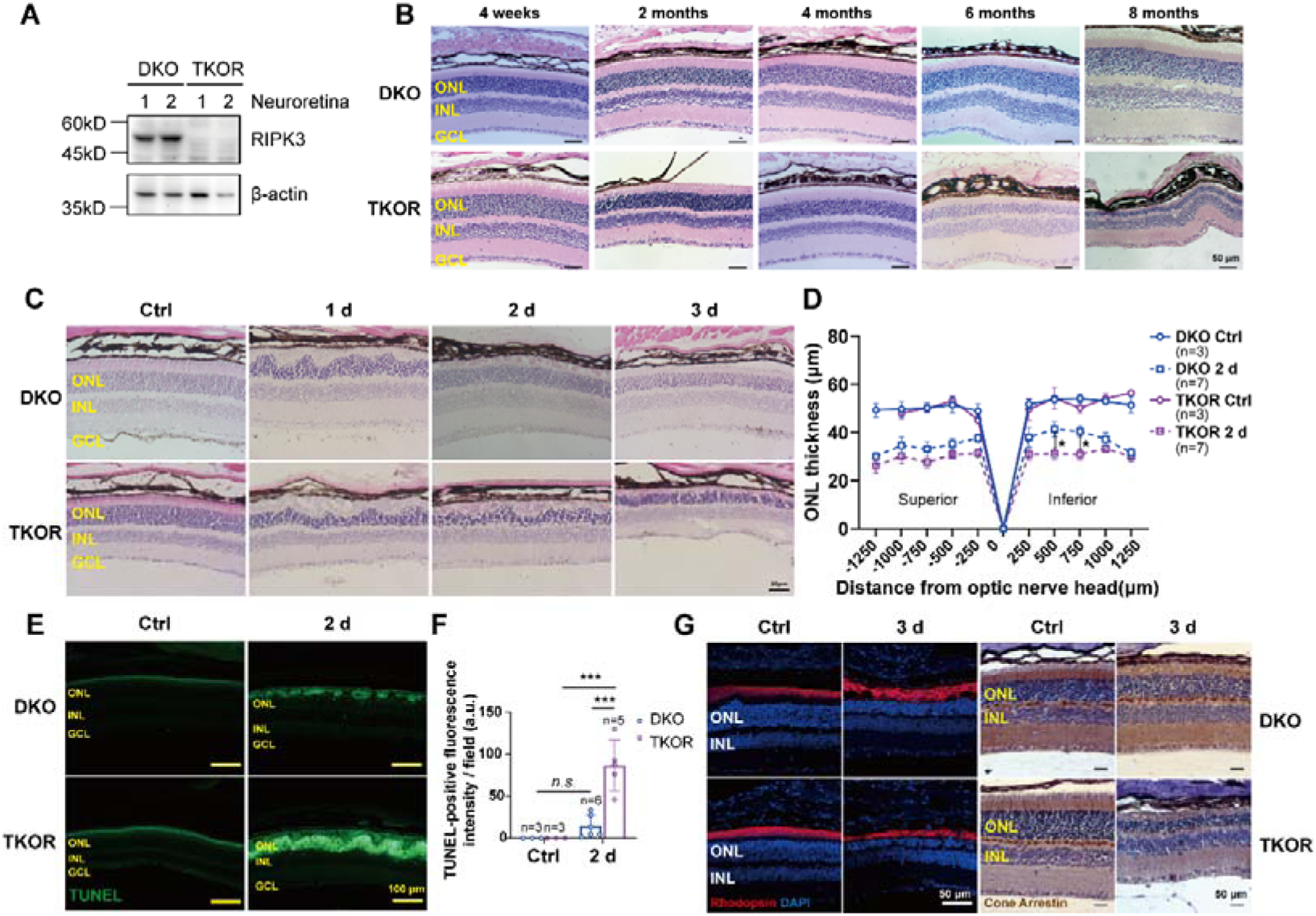
Knockout of *Ripk3* accelerates photoreceptor degeneration in STGD1 mouse model. (**A**) Western blot analysis of RIPK3 in the neuroretina of DKO and abca4^-/-^ rdh8^-/-^ *ripk3^-/-^* (TKOR) mice. β-actin was used as a loading control. (**B**) Representative hematoxylin and eosin (H&E) staining images of mouse retinas from DKO and TKOR mice at different ages under normal light (Scale bar = 50 μm). (**C**) Representative hematoxylin and eosin (H&E) staining images of retinas from DKO and TKOR mice before or at various time points after a 30-min exposure to 10,000 lux white LED light (Scale bar = 50 μm). (**D**) Quantification of ONL thickness in DKO and TKOR mice before and at 2 d after light exposure. Data are obtained from 3∼7 mice per group. (**E**) Representative terminal deoxynucleotidyl transferase dUTP nick end labeling (TUNEL) staining images of retinal cross-sections from DKO and TKOR mice before or at 2 d after light exposure (Scale bar = 100 μm). (**F**) Quantification of TUNEL fluorescence intensity in the ONL of DKO and TKOR mice before or at 2 d after light exposure. Data are obtained from 3∼6 mice per group. (**G**) Representative images of immunofluorescence staining of rod rhodopsin or immunohistochemistry staining of cone arrestin in retinal cross-sections from DKO and TKOR mice before or at 2 d after light exposure (Scale bar = 50 μm). ONL: outer nuclear layer; INL: inner nuclear layer; GCL: ganglion cell layer. For all the experiments, statistical significance was determined by two-way ANOVA with Tukey’s post-hoc test: \**p* < 0.05, \*\**p* < 0.01, \*\*\**p* < 0.001, *n.s.*: not significant. All data are shown as the mean ± standard deviation (S.D.).

To identify the mode of photoreceptor cell death, propidium iodide (PI) was injected into the vitreous body to assess photoreceptor cell membrane permeability at 3 d after light exposure when most of photoreceptors were positive for TUNEL staining in both DKO and TKOR mice. Nuclear PI staining was barely detectable in the ONL of both DKO and TKOR mice (**Figures** 5**A** and **B**), suggesting that light exposure did not compromise photoreceptor plasma membrane integrity, regardless of *Ripk3* status. Furthermore, the ultrastructural morphology of photoreceptors was characterized by transmission electron microscopy (TEM). Under TEM, most photoreceptors displaying chromatin condensation were identified as apoptotic cells, whereas those characterized by cytoplasmic vacuolization were defined as necrotic cells (**Figure** 5**C**)^(22, 44)^.

**Figure 5.**
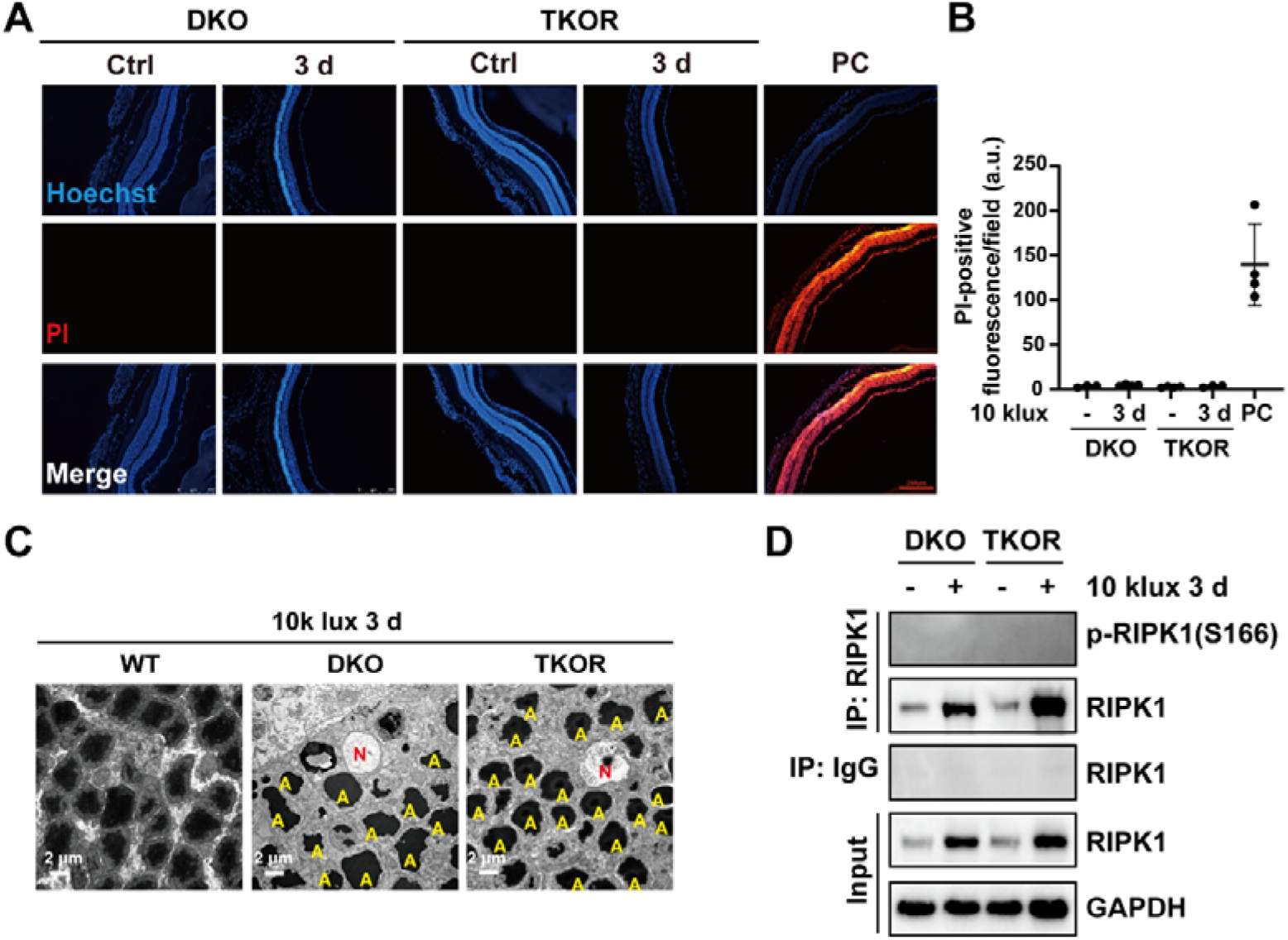
Apoptosis is the predominant mode of photoreceptor cell death in STGD1 mouse model after light exposure. (**A**) Representative immunofluorescence images of propidium iodide (PI) staining after intravitreal injection in the neuroretina of DKO and TKOR mice before or at 3 d after 30-min exposure to 10,000 lux full-spectrum white light (Scale bar = 250 μm). PC: positive control, in which retinal cross-sections were permeabilized with 0.2% Triton X-100 and stained with PI. (**B**) Quantification of PI-positive fluorescence intensities in the retinal cross-sections of DKO and TKOR mice. Data were collected from 3∼4 mice per group. (**C**) Representative transmission electron microscope (TEM) images of the ONL in the central retinas of WT, DKO and TKOR mice at 3 d after 30-min exposure to white LED light (Scale bar = 2 μm). A: Apoptosis; N: Necrosis. (**D**) Immunoprecipitation analysis of RIPK1 and its phosphorylation modifications in the retinas of DKO and TKOR mice before or at 3 d after light exposure. GAPDH was used as a loading control.

In addition to apoptosis, parthanatos, a distinct form of cell death mediated by poly(ADP-ribose) polymerase 1 (PARP-1) and apoptosis-inducing factor (AIF), is also characterized by nuclear condensation (45). In response to DNA damage, PARP-1 enzymatic activity is upregulated, leading to rapid polymerization of poly (ADP-ribose) (PAR) chains; the accumulation of these chains is closely associated with the initiation of cell death (46). Although PAR modification was enhanced in the retinas of both DKO and TKOR mice after light exposure, its level was significantly lower in TKOR mice than in DKO mice at 3 d after light exposure (**Figure** S3**A**). Downstream of PARP-1 activation, AIF is cleaved by calpains and/or cathepsins to generate truncated AIF (tAIF), which promotes tAIF translocation from mitochondria to the nucleus (47). The nuclear tAIF induces chromatinolysis, and thus ultimately triggers cell death (47). Notably, however, tAIF was undetectable in the retinas of either mouse strain following light exposure (**Figure** S3**A**). Immunohistochemical analysis of retinal cross-sections revealed that the most robust AIF staining was localized to the mitochondrial-dense inner segments of photoreceptors in the control groups of both DKO and TKOR mice (**Figure** S3**B**). Despite substantial photoreceptor loss triggered by light exposure, there was no evident nuclear translocation of AIF in these cells (**Figure** S3**B**). Collectively, these results indicate that parthanatos does not serve as a major mode of photoreceptor cell death modulated by RIPK3 following light exposure. RIPK1 is a critical regulator of multiple cell death pathways downstream of TNFα, including RIPK1-independent apoptosis, necroptosis, and RIPK1-dependent apoptosis (RDA) (38). The latter two pathways are characterized by phosphorylation of RIPK1 on Serine 166 (38). To further confirm whether apoptosis or necroptosis was involved in photoreceptor cell death after light exposure in DKO and TKOR mice, immunoprecipitation assays were performed to enrich RIPK1 from retinal lysates at 3 d after light exposure when most of photoreceptor cells were undergoing cell death. Although the protein levels of RIPK1 were increased in the retinas of both DKO and TKOR mice, phosphorylation of RIPK1 at Ser166, a key post-translational modification indicative of RDA activation and necroptosis, remained undetectable (**Figure** 5**D**). This suggests that neither necroptosis nor RDA contributed significantly to retinal degeneration in these models.

Taken together, these results indicate that the majority of photoreceptors undergo apoptosis after light exposure in both DKO and TKOR mice.

### RIPK3 Suppresses Extrinsic Apoptosis in the Retinas of DKO Mice and atRAL-treated Cells

To further elucidate the regulatory role of RIPK3, neuroretina tissues from DKO and TKOR mice at 2 d after light exposure were subjected to bulk RNA-seq analysis. Using 2-day-light-exposed neuroretinas from DKO mice as the control cohort, a total of 2,236 DEGs were identified (*p-*value < 0.05, folds of change ≥ 1.2), comprising 1249 upregulated and 987 downregulated transcripts. KEGG pathway enrichment analysis demonstrated that these DEGs were also significantly enriched in the TNF signaling pathway(mmu04668) and Apoptosis pathway(mmu04210) (**Figure** 6**A**). Moreover, a heatmap visualization of the identified DEGs associated with the apoptosis pathway showed that genes encoding death ligands and receptors, including *Tnf*, *Tnfsf10*, *Tnfrsf1a*, and *Fas*, along with key genes in the extrinsic apoptosis cascade, such as *Casp8* and *Casp3,* were upregulated in the TKOR retinas compared with DKO controls at 2 d after light exposure (**Figure** 6**B**).Additionally, qRT-PCR analysis confirmed that the mRNA levels of *Casp8* and *Casp3* were significantly higher in the retinas of TKOR mice compared with those of DKO mice at 2 d post light exposure (**Figure** 6**C**, *Casp8*: DKO vs. TKOR, 0d: *p* > 0.9999; 1d: *p* > 0.9999; 2d: *p* = 0.0006; 3d: 0.9999; *Casp3*: DKO vs. TKOR, 0d: *p* > 0.9999; 1d: *p* = 0.9998; 2d: *p* = 0.0210; 3d: *p* > 0.9999).

**Figure 6.**
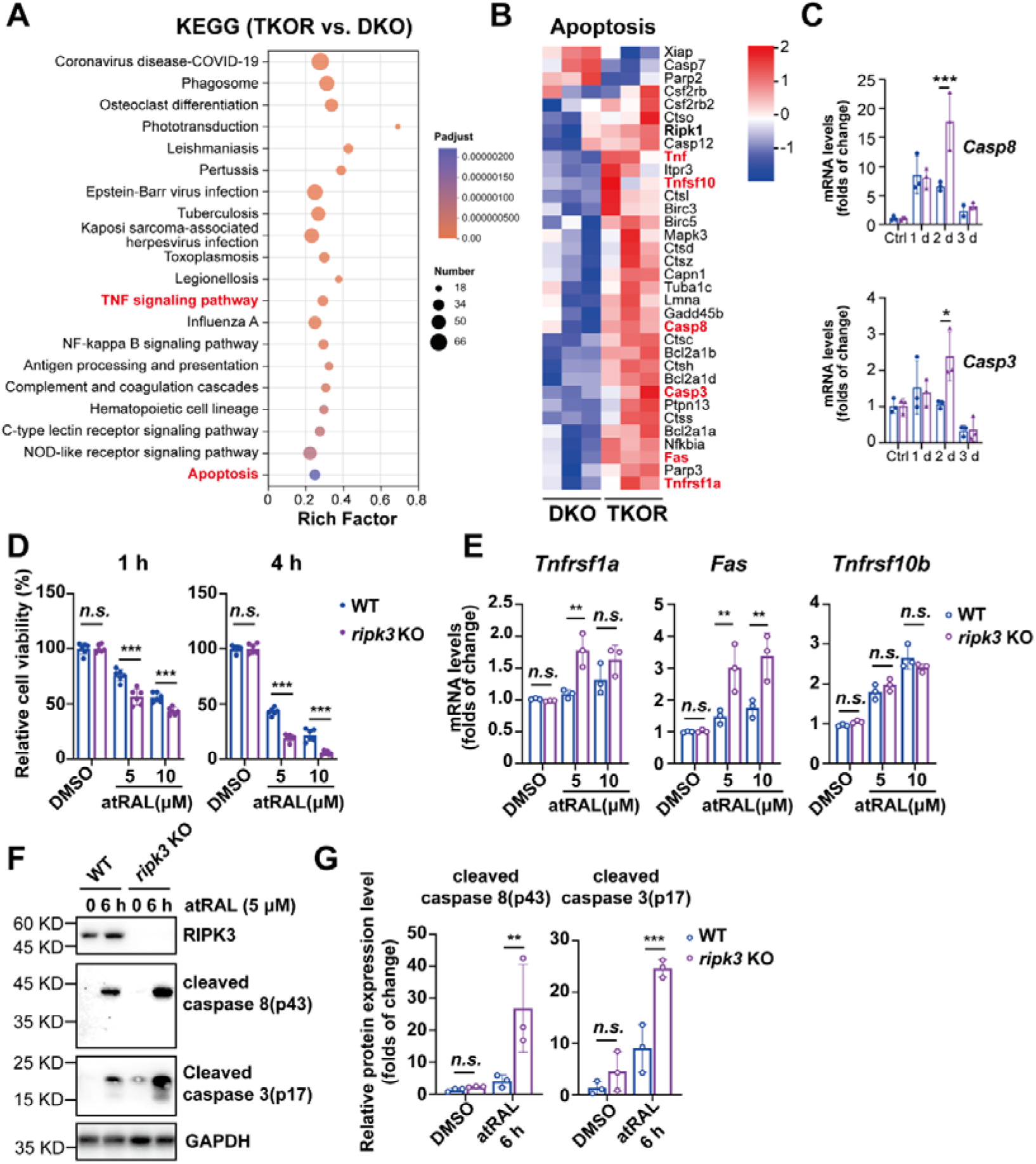
Loss of RIPK3 enhances extrinsic apoptosis in vivo and in vitro. (**A**) KEGG pathway enrichment analysis of DEGs in the neuroretinas of DKO and TKOR mice at 2 d after a 30-min exposure to white LED light. (**B**) Heatmap of DEGs in the apoptosis pathway identified by RNA sequencing in the neuroretinas of DKO and TKOR mice. (**C**) Quantitative RT-PCR analysis of *Casp8* and *Casp3* mRNA expression levels in the retinas of DKO and TKOR mice before or after a 30-min exposure to white LED light. Data were obtained from 3 mice per group. (**D**) WT and *ripk3* KO 661W cells were incubated with 5 or 10 μM of atRAL, and cell viability was evaluated by CCK8 assay. (**E**) Quantitative RT-PCR analysis of *Tnfrsf1a, Fas,* and *Tnfrsf10b* mRNA expression levels in WT and *ripk3* KO 661W cells at 6 h after atRAL treatment. Data were obtained from 3 independent experiments. (**F**) Representative Western blot images showing cleaved caspase 8 (p43) and cleaved caspase 3(p17) in WT and *ripk3* KO 661W cell lysates following treatment with 5 μM atRAL for 6 h. GAPDH was used as a loading control. (**G**) Quantification results of the protein levels of cleaved caspase 8 (p43) and cleaved caspase 3 (p17) in (**F**). For all the experiments, statistical significance was determined by two-way ANOVA with Tukey’s *post-hoc* test: \**p* < 0.05, \*\**p* < 0.01, \*\*\**p* < 0.001. All data are shown as the mean ± standard deviation (S.D.).

To further validate that RIPK3 modulates extrinsic apoptosis, we established a *Ripk3* knockout (*ripk3* KO) 661W cell line. Cell viability assays demonstrated that *Ripk3* deficiency significantly increased the vulnerability of 661W cells to atRAL-induced cytotoxicity (**Figure** 6**D**, WT vs. *ripk3* KO, 1h: DMSO, *p* > 0.9999; 5 μM, *p* < 0.0001; 10 μM, *p* = 0.0003; 4h: DMSO, *p* > 0.9999;5 μM, *p* < 0.0001; 10 μM, *p* < 0.0001). Moreover, qRT-PCR analysis revealed that atRAL treatment upregulated the transcription of multiple cell death receptor genes, including *Tnfrsf1a*, *Fas*, and *Tnfrsf10b* (**Figure** 6**E**), which was consistent with the mouse RNA-seq results. Notably, the mRNA expression levels of *Tnfrsf1a* and *Fas* were further elevated in *ripk3* KO cells relative to WT cells following atRAL exposure (**Figure** 6**E**, WT vs. *ripk3* KO, *Tnfrsf1a*: DMSO, *p* = 0.9951; 5 μM atRAL, *p* = 0.0013; 10 μM atRAL, *p* = 0.1355; *Fas*: DMSO, *p*=0.9999; 5 μM atRAL, *p* = 0.0038; 10 μM atRAL, *p* = 0.0024). In line with these transcriptional changes, Western blot assays showed that the levels of cleaved caspase 8(p43) and cleaved caspase 3(p17) were significantly higher in *ripk3* KO cells than in WT cells after atRAL treatment (**Figures** 6**F** and **G**, WT vs. *ripk3* KO, caspase 8(p43): DMSO, *p* = 0.9830; 6 h, *p* = 0.0080; caspase 3(p17): DMSO, *p* = 0.4438; 6 h, *p* = 0.0007). These results confirm that RIPK3 acts as a negative regulator of extrinsic apoptosis in photoreceptor cells in STGD1 models.

### Pharmacological Inhibition of Caspases Partially Rescues Photoreceptors in TKOR Mice

To determine whether caspase-dependent apoptosis contributes to photoreceptor degeneration in STGD1 mouse models, we administered the pan-caspase inhibitor Z-VAD-FMK via intravitreal injection following light exposure. In TKOR mice, Z-VAD-FMK treatment significantly attenuated photoreceptor cell loss compared to vehicle-treated controls (**Figures** 7**A** and **B**; TKOR Ctrl vs. TKOR 5 d: *p*L<L0.0001; TKOR 5 d vs. TKOR 5 d + Z-VAD-FMK: *p*L=L0.0337). The thickness of the outer nuclear layer (ONL) in Z-VAD-FMK–treated TKOR retinas was comparable to that observed in light-exposed DKO mice without caspase inhibition. In contrast, inhibition of caspases in DKO mice exacerbated light-induced photoreceptor cell death (**Figures** 7**A** and **B**). In addition, intravitreal injection of caspase-8-specific inhibitor Z-IETD-FMK conferred a level of photoreceptor protection indistinguishable from that achieved with Z-VAD-FMK in TKOR mice (**Figure** S4). Taken together, our data support a model in which RIPK3 suppresses caspase-8–mediated extrinsic apoptosis in STGD1 retinas, thereby exerting a protective role in photoreceptor degeneration.

**Figure 7.**
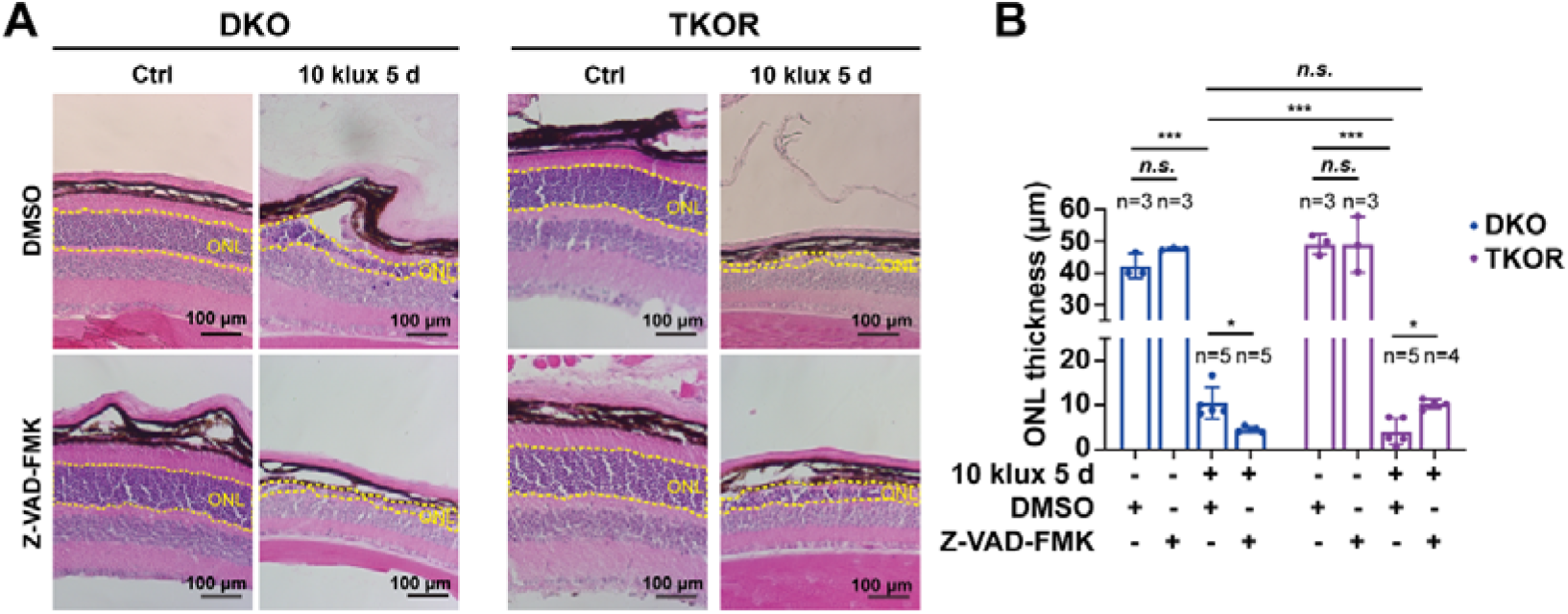
Pharmacological inhibition of caspases partially rescues photoreceptors in TKOR mice. (**A**) Representative hematoxylin and eosin (H&E) staining images of mouse retinas from DKO and TKOR mice before or at 5 d after a 30-min exposure to 10,000 lux white LED light, with or without intravitreal injections of Z-VAD-FMK (Scale bar = 100 μm). ONL: outer nuclear layer. (**B**) Quantification of central retinal ONL thickness in DKO and TKOR mice before or at 5 d after light exposure, with or without intravitreal injections of Z-VAD-FMK. Data are obtained from 3∼5 mice per group. For all the experiments, statistical significance was determined by two-way ANOVA with Tukey’s *post-hoc* test: \**p* < 0.05, \*\**p* < 0.01, \*\*\**p* < 0.001. All data are shown as the mean ± standard deviation (S.D.).

## Discussion

The most interesting finding of the present study is that RIPK3 acts as a suppressor of extrinsic apoptosis rather than a mediator of necroptosis in the retinas of STGD1 models. While RIPK3 is well characterized as a core regulator of necroptosis, its involvement in apoptotic pathways has been documented in previous studies, albeit with conflicting conclusions. Early in vitro studies demonstrated that RIPK3 exerts a pro-apoptotic effect when overexpressed in multiple cell lines (19, 48, 49). Moreover, in vivo evidence showed that genetically engineered mice harboring the kinase-dead RIPK3 D161N mutant die around embryonic day 11.5 due to increased RIPK1/caspase-8-dependent apoptosis in endothelial cells (21), suggesting that the kinase activity of RIPK3 is indispensable for maintaining cell survival. In contrast, a subsequent study utilizing RIPK3 kinase inhibitors and a panel of different kinase-dead mutants revealed that the pro-apoptotic function of RIPK3 is independent of its catalytic activity, but instead relies on its RIP homotypic interaction motif (RHIM) domain to recruit RIPK1 and activate caspase-8 (20). These conflicting reports indicate that while RIPK3 is known to modulate RIPK1/caspase-8-mediated extrinsic apoptosis, the precise mechanisms underlying its functional divergence in this pathway remain largely elusive. In this study, we found that RIPK3 was upregulated in the retinas of DKO mice (**Figures** 3**A** and **B**); however, this upregulation exerted a protective role by suppressing extrinsic apoptosis in photoreceptors (**Figures** 5 and 6). Consistent with our findings, RIPK3 has been shown to inhibit hepatocyte apoptosis in a high-fat diet (HFD)-induced liver injury model ^49^. Integrating both in vivo and in vitro evidence from the literature and our own work, we speculate that RIPK3 is not a unidirectional death effector but a context-dependent signaling hub whose functional output, either pro-death or pro-survival, is shaped by cellular identity, nature of stress, expression level, conformational state, and possibly post-translational modifications. Necroptosis is a form of regulated necrosis that is activated when apoptosis, particularly caspase-8-mediated extrinsic apoptosis, is inhibited. Downstream of death receptors, pattern recognition receptors (e.g., Toll-like receptor (TLR)-3 and -4) or zDNA-binding protein 1 (ZBP1), RIPK3 undergoes phosphorylation when necroptosis is activated; this in turn phosphorylates MLKL to execute cell death (50, 51). Therefore, MLKL is regarded as the executor of necroptosis. In the present work, multiple lines of evidence indicate that necroptosis does not contribute directly to photoreceptor degeneration in DKO mice. Biochemical analysis revealed no MLKL phosphorylation (**Figure** 3**A**), and genetic ablation of *Mlkl* failed to rescue photoreceptor loss (**Figures** 3**E**-**H**). Consistent with these in vivo observations, exposure of 661W cells to atRAL did not induce MLKL phosphorylation and other hallmarks of necroptosis (**Figures** 3**C** and **D**). All these findings align with earlier reports by Cai et al (31), who also failed to detect MLKL phosphorylation in DKO mice after light exposure. In contrast, another study reported MLKL phosphorylation in the neuroretina after mice were exposed to 15,000 lux light for 2 h in both wildtype 129S2/Sv and RPE65-null rd12 mouse models (26). One possible explanation for the distinct results between our study and previous reports might be due to the differences of light intensities as we only used a relatively mild regimen—10,000 lux light to illuminate mice for 30 min. Under such conditions, photoreceptor cells may preferentially initiate the apoptosis program, while necroptosis remains inactive. Supporting this interpretation, under our condition light exposure did not compromise the integrity of the photoreceptor plasma membrane regardless of *Ripk3* status (**Figures** 5**A** and **B**), further confirming the absence of necroptosis in our model. We propose that the degree of retinal damage dictates the dominant type of photoreceptor cell death. Notably, we found that photoreceptor degeneration in DKO mice was aggravated when apoptosis was inhibited by either the pan-caspase inhibitor Z-VAD-FMK (**Figure** 7) or the caspase 8 inhibitor Z-IETD-FMK (**Figure** S4). This indicates that blocking apoptosis may shift the mode of cell death toward necroptosis, a phenomenon previously described in a retinal detachment model (22). Moreover, while we observed accelerated degeneration of both rod and cone photoreceptor cells in *Ripk3*-deficient mice (**Figure** 4**G**), other studies have shown that RIPK3 mediates necrotic cone cell death in the rd10 mouse model (30). Given all these seemingly contradictory findings, further mechanistic studies are warranted to clarify the precise conditions under which RIPK3 promotes, suppresses, or switches between distinct cell death modalities in the retina.

KEGG analysis of DEGs revealed that TNF signaling pathway was activated in the retinas of DKO mice compared with WT mice after light exposure (**Figure** 1). Moreover, the activation level of this pathway was further elevated in the retinas of light-exposed TKOR mice compared to DKO mice (**Figure** 6). Downstream of TNF receptor 1 (TNFR1), in-depth dissection of biochemical pathways has uncovered the complex crosstalk among different cell death modalities. Specifically, blockage of caspase (executor of apoptosis) activity triggers necroptosis, whereas overactivation of caspases cleaves gasdermins to activate pyroptosis (52, 53). Conversely, cleavage of RIPK3 by caspase 8 restricts pyroptosis dependent on NLRP3 inflammasome activation (54). The results in this study indicated that apoptosis was the predominant type of cell death in the retinas of both DKO and TKOR mice after light exposure (**Figure** 5). However, intravitreal injection of caspase inhibitors only partially protected photoreceptors in TKOR mice (**Figures** 7 and S4). One potential explanation for this observation is that caspase inhibitors were rapidly metabolized in the mouse vitreous, and thus were failing to fully suppress caspase activation. Nevertheless, we could not rule out the possibility that other modes of cell death may be triggered during photoreceptor degeneration.

Downstream of TNF signaling, RIPK1 acts as a critical regulator of both apoptosis and necroptosis ^(38)^. Activation of TNFR1 recruits a transient multimeric complex (complex I) containing RIPK1, which promotes the activation of the pro-survival nuclear factor-κB (NF-κB) pathway and inhibits RIPK1 kinase activation through complex ubiquitination and phosphorylation events. When protein synthesis is inhibited, TNF can induce RIPK1-independent apoptosis, in which caspase 8 cleaves RIPK1 at Asp324 to suppress necroptosis. Impairment of RIPK1 ubiquitination or inhibitory phosphorylation leads to phosphorylation of RIPK1 at Ser166, which drives the formation of a RIPK1-TRADD-FADD-caspase 8 complex (complex IIa), thereby triggering RIPK1-dependent apoptosis (RDA). Additionally, when caspases are inhibited or caspase 8 is deficient, Ser166-phosphorylated RIPK1 at Ser166 interacts with RIPK3 via their RHIM domains. Subsequently, the formation of RIPK1-RIPK3-MLKL complex (complex IIb) initiates necroptosis ^(38)^. In this study, we also found that the protein levels of full-length RIPK1 were elevated in the retinas of both DKO and TKOR mice after light exposure (**Figure** 5**D**). However, the phosphorylation of RIPK1 at Ser166 remained undetectable, even after RIPK1 enrichment by immunoprecipitation (**Figure** 5**D**). These results further support our conclusion that necroptosis is not involved in photoreceptor degeneration in DKO mice. Nevertheless, the extent to which RIPK1 participates in the regulatory network governing photoreceptor degeneration in this STGD1 mouse model remains unclear and it will be investigated in future studies.

Beyond apoptosis and necroptosis, we also tried to investigate whether parthanatos contributes to photoreceptor cell death in STGD1. One piece of evidence supporting the induction of parthanatos is the increased PAR modifications in the retinas of both DKO and TKOR mice following light exposure (**Figure** S3**A**). However, we failed to detect AIF truncation and its nuclear translocation (**Figure** S3), which are both hallmarks of parthanatos (55). Moreover, parthanatos is caspase-independent, whereas our results suggested that photoreceptor degeneration in our model was at least partially caspase-dependent (**Figures** 7 and S4). Collectively, we conclude that parthanatos does not serve as a major form of cell death in STGD1. Additionally, although RIPK3 deficiency accelerated photoreceptor cell death, the levels of PAR modification were lower in the retinas of TKOR mice compared to DKO mice (**Figure** S3), which suggests that RIPK3 does not regulate photoreceptor degeneration in STGD1 by modulating the levels of parthanatos.

This study has several limitations. First, the majority of our experiments were conducted in an acute light-induced retinal injury model, which does not fully recapitulate the relatively slow, progressive photoreceptor degeneration characteristic of STGD1. Nevertheless, we also observed that *Ripk3* deficiency exacerbated age-dependent photoreceptor loss in DKO mice (**Figure** 4**B**), suggesting a conserved role of RIPK3 across different degenerative contexts. Second, our in vitro work relied on the 661W cell line, a cone-like line derived from retinal tumors of transgenic mice expressing SV40 large T antigen under the control of the human IRBP promoter (35). While 661W cells express several key marker genes of cone photoreceptors, they differ from native cells in several aspects, including continuous proliferation, loss of cellular polarity, and absence of outer segments (35, 56). Despite these caveats, the concordant protective function of RIPK3 observed both in vivo and in 661W cells supports the utility of this cell line as a model for mechanistic exploration in our experimental context.

In conclusion, our study uncovers a regulatory role of RIPK3 in photoreceptor apoptosis, findings that likely only reveals the tip of the iceberg of the intricate cell death regulatory network governing photoreceptor degeneration. Notably, as RIPK3 functions as a suppressor of photoreceptor apoptosis, how to modulate RIPK3 so as to protect photoreceptors in STGD1 remains as an open question. Accordingly, to deepen our understanding of the regulatory mechanisms underlying photoreceptor cell death and ultimately develop effective photoreceptor-protective strategies for STGD1 and potentially AMD, several key questions must be addressed in future studies: how RIPK3 switches its functional roles across distinct cell death modalities, the precise contribution of RIPK1 to this regulatory network, and the cooperative manner in which different cell death pathways orchestrate photoreceptor degeneration.

## Acknowledgement

This work was supported by grants from the Natural Science Foundation of Xiamen, China (Grant No.3502Z202373024), the Natural Science Foundation of Xiamen (Grant No.3502Z20227099), the Natural Science Foundation of Fujian Province (Grant No.2022J05297), and China National Natural Science Foundation (Grant No. 81700864). The authors thank Prof. Juan Lin from School of Medicine, Xiamen University and Prof. Steven J. Fliesler from University at Buffalo, The State University of New York (SUNY) for insightful suggestions, Jingru Huang and Xiang You from Central Lab, School of Medicine, Xiamen University for technical support in confocal imaging.

## Data Availability

All data supporting the findings of this study are available from the corresponding author upon reasonable request.

